# Eye movements reveal the intuitive understanding of materials and their behavior

**DOI:** 10.64898/2026.01.19.700384

**Authors:** Alexander Goettker, Filipp Schmidt

## Abstract

Does a metal ball sink or float when dropped into water? Answering this intuitively simple question relies on our understanding of materials and their physics^1^. Knowing materials helps us to recognize objects, estimate their properties, and predict how they will behave when interacting with other objects or materials^2,3^. This leads to expectations about material behaviors already in young infants^4,5^, for example, that a metal ball should sink in water. Typically, properties and affordances of materials have been studied with perceptual tasks. But even though explicit perceptual judgments allow us to identify important determinants of material understanding, it is not clear how they relate to dynamic predictions of material behavior — which we need for successful interactions with objects^3^. Here, we tap into material understanding by leveraging anticipatory eye movements, which have been shown to be direct readouts of trajectory predictions^6^ and can also reflect expectations about material properties such as elasticity or mass^7^. This allows us to reveal the temporal evolution of dynamic predictions based on material properties. Furthermore, by using novel stimuli that violate intuitive expectations about material behavior, we demonstrate that even though dynamic predictions can be learned over time, there remains a fundamental difference between predictions in line with and against our intuitive expectations.

## Main Text

To study material understanding through anticipatory eye movements, we created a novel set of short animations in which balls made from either metal or plastic dropped into a pool of water (**Fig. 1A**). To vary expectations and increase stimulus diversity, we rendered the balls with five different plastics (e.g., beach ball, bubble wrap) and five metals (e.g., iron, copper) (**Fig. 1B**). Crucially, for the first 2 s of each animation, all balls followed the same visual motion path. Only after contact with the water did their behaviors diverge: some balls sank rapidly, while some balls bobbed and floated. Thirty naïve participants saw the full animations and had to press a button when the ball entered a target area (see **Fig. 1A**). Critically, we used three different conditions: In the **Consistent** condition, the ball behaved as intuitively expected (metal balls sank, plastic balls floated). In the **Inconsistent** condition, the behavior of the balls was opposite to the intuitive expectation (metal balls floated, plastic balls sank), but still completely predictable (**Fig. 1C**). To balance the experience people had with our animations, the order of the Consistent and Inconsistent condition was varied. For comparison, we also included a **Mask** condition, where the ball material was concealed (uniform black) and its behavior unpredictable.

**Figure 1.**
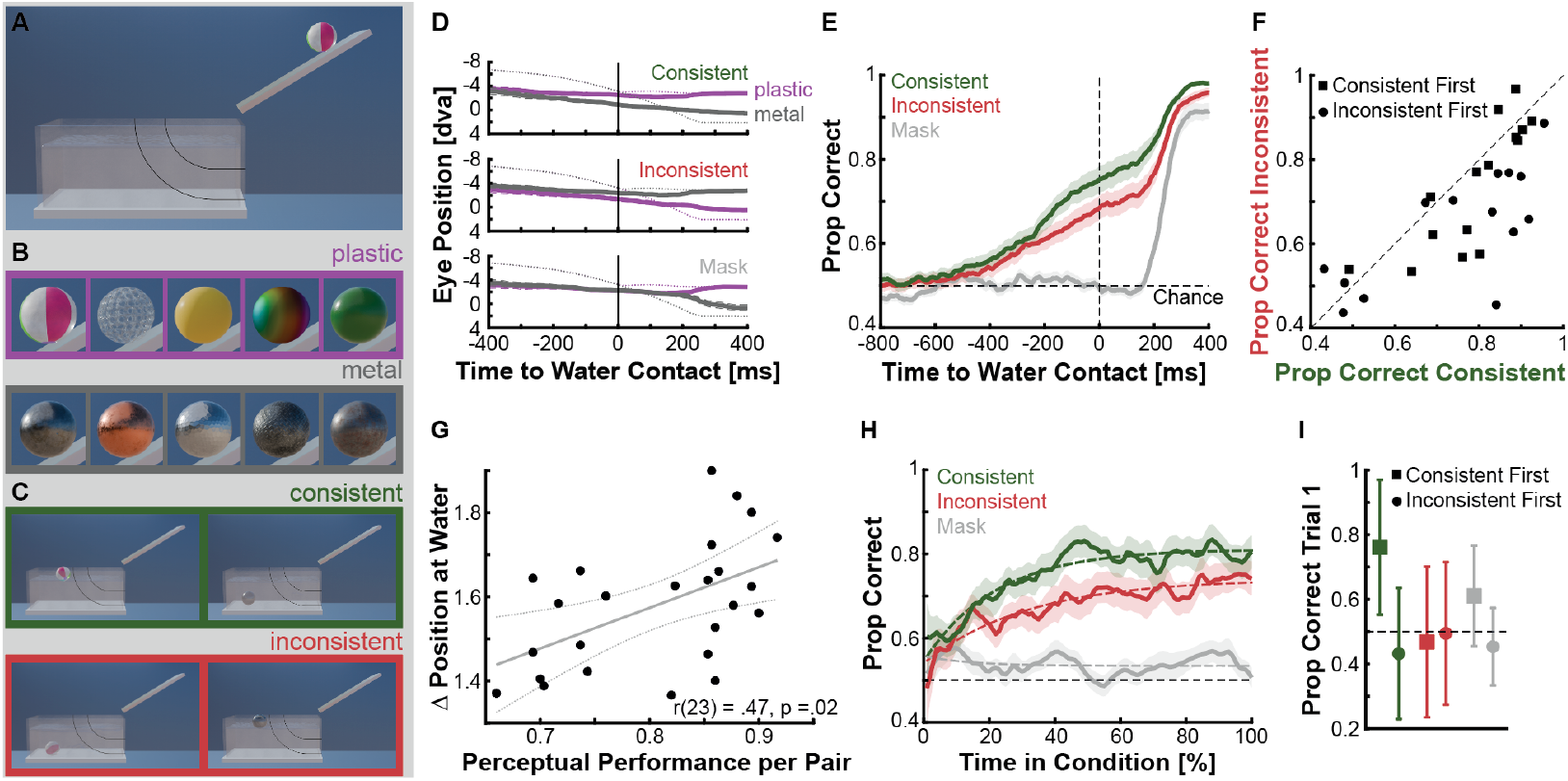
Dynamic Gaze predictions based on material properties. (**A**) Example stimulus, showing the first frame of an animation with a plastic ball on the ramp and the target area (black lines). (**B**) All plastic (purple) and metal (grey) materials. (**C**) Example final animation frames for the Consistent (green: plastic floating, metal sinking) and Inconsistent (red: plastic sinking, metal floating) conditions. (**D**) Vertical eye positions as a function of time relative to water contact, for both materials and the three conditions (subplots). (**E**) Corresponding classification accuracy within a trial as a function of time. (**F**) Comparison of classification accuracy at water contact for the Consistent and Inconsistent condition. (**g**) Relationship between perception of the materials and vertical eye position at water contact. (**h**) Classification accuracy as a function of visual experience in each condition. (**I**) Classification accuracy in the first trial. Different colors indicate the different materials or conditions. All shaded areas depict the standard error. Error bars show the 95% CI.

When looking at the average vertical gaze traces relative to the moment of water contact (**Fig. 1D**), we found that the oculomotor system reflects a dynamic prediction of future stimulus behavior: Eye movements diverged *before* the physical divergence of the two possibilities (sinking vs. floating) in the Consistent and Inconsistent conditions, but only *after* the water contact in the Mask condition. To quantify this effect, we trained a classifier to predict whether the ball will be floating or sinking from the continuous gaze traces (**Fig. 1E**). There are two key results. First, above-chance classification performance for Consistent and Inconsistent conditions emerges from around 400 ms *before* the physical change of behavior at the moment of water contact.

Thus, in the two predictable conditions, the eyes successfully anticipated and reflected the future trajectories. In contrast, in the Mask condition, this only happens about 200 ms after water contact, demonstrating purely reactive behavior. Second, although Consistent and Inconsistent conditions were both fully predictable, anticipatory eye movements were different: average classification performance at the moment of the water contact was significantly higher for the Consistent condition (**Fig. 1F**, t(29) = 3.256, p = .003). This indicates that predictions that align with our sense of intuitive physics are fundamentally different from more recently learned relationships. The unique nature of these predictions is supported by two additional observations.

First, we observed that the differences in eye movement behavior scaled with the uncertainty about material properties. Before the main experiment, all participants were shown the initial phase of our animations until right before the moment of water contact. Then they judged for each of the materials (5 plastics, 5 metals), whether the ball would float or sink. We then computed for each possible pair of metallic and plastic stimuli the average accuracy of these judgments, and related it to the average vertical difference in gaze position for the two stimuli at water contact. Across all pairs, we found a significant relationship (r(23) = .469, p = .018, see **Fig. 1G**), indicating that stronger expectations about materials are related to stronger effects in eye movement behavior.

Second, we observed a stable benefit for the Consistent condition across the whole experiment. We again used a classification approach to quantify the development of the predictions over time within each condition. We trained the classifier with the gaze position at water contact from a random selection of trials, and then predicted each trial individually to compare performance over time (see Methods for more details). The results show a constant offset between the Consistent and Inconsistent condition (**Fig. 1H**), with lower prediction levels and a lower asymptote in the Inconsistent condition (Bootstrap estimates, Consistent: Median = 0.806, 95%CI = 0.787-0.842, Inconsistent: Median = 0.739, 95%CI = 0.712-0.803) even after learning (i.e., in the last trial). Additionally, behavior was already different in the very first trial (**Fig. 1I**): Performance was only significantly above chance for participants seeing the Consistent condition first. In all other conditions — also when seeing the Inconsistent animations first and thus being primed for an expectation violation — people were not able to successfully anticipate ball behavior from the outset.

Together, our results show that eye movements can serve as an objective, implicit, and rich marker for our intuitive understanding of materials and their behavior. Simply put, what we think an object *is made of* shapes predictions about what it *will do* — and our eyes provide a sensitive window into this process as it unfolds in real time. This anticipatory integration of material knowledge extends previous findings on context-driven predictions^6^ and shape-driven anticipatory smooth pursuit^8^ to the domain of physical reasoning. Our knowledge about materials is arguably one of the earliest-developing forms of physical understanding^4,5^ and fundamental for object handling, including tool use, foraging, and navigation in natural environments^2,3^. That eye movements align gaze with the expected, not yet observed, trajectory of the object, minimizes the negative consequences of latencies of visual and motor processing and helps to prepare future interactions with objects^9^.

In contrast to classic material perception studies that rely on explicit perceptual judgements, we leveraged anticipatory eye movements to reveal three novel features of dynamic predictions about materials. First, we show that predictions start to shape our behavior from around 400 ms before the actual event **(Fig 1e)**. Second, these dynamic predictions reflect perceptual uncertainty about the stimulus: They are stronger for objects with certainly known material properties (**Fig. 1g**). Third, predictions that match our intuitive expectations are special^10^: While our participants also learned to anticipate movements in the Inconsistent condition, for intuitively correct stimuli predictions were stronger, already present from the first trial, and stayed more pronounced throughout the experiment. These insights on dynamic predictions based on material understanding also open new avenues for studying the neural computations underlying intuitive physics. Where do differences between Consistent and Inconsistent conditions emerge? Are intuitive predictions really unique, or is there a level of exposure that can make predictions in the Inconsistent condition similarly strong? Together, our results highlight that expectations about materials and our intuitive understanding of the world are not confined to high-level cognition but are woven into the earliest stages of sensorimotor planning^10^.

## Acknowledgement

The authors want to thank Svea Kürthen for her help with data collection. This work was funded by the Deutsche Forschungsgemeinschaft (German Research Foundation, DFG, project number 222641018–SFB/TRR 135 Projects A1 and C1).

## Data Availability

All Data, and Stimuli are available at OSF: https://osf.io/e39hp. Analysis Code is on Github: https://github.com/AlexanderGoettker/MaterialAnticipation

## Supplemental Information

Supplemental Information includes a detailed description of the methods, a link to the publicly available data set and experimental and analysis code.

## Author Contributions

Conceptualization & Methodology, A.G, F.S.; Software and Formal Analysis, A.G.; Writing – Original Draft, A.G., F.S.; Writing – Review & Editing, A.G., F.S.; Visualization, A.G., F.S.; Supervision and Funding Acquisition, A.G., F.S.

## Methods

### Participants

30 human observers (mean age 24.7 years; standard deviation: 4.4 years; 25 female) completed the tasks. All participants reported normal or corrected-to-normal vision and were naïve with respect to the study. Experimental procedures were in line with the Declaration of Helsinki and were approved by the Local Ethics Committee of the Department of Psychology and Sports Sciences of the Justus Liebig University Giessen (LEK-2021-0032). Written informed consent was obtained from each participant. Participants received 8 Euros compensation per hour or course credits.

### Experimental Setup

Participants sat at a table in a dimly illuminated room with their heads positioned in a combined chin and forehead rest. Their eyes were roughly aligned with the upper half of a monitor (60 cm × 32 cm, 3840 × 2160 pixels; Phillips, Amsterdam, Netherlands) at a distance of 70 cm. Under these conditions, the monitor spanned approximately 49 × 26 dva. The experiment was programmed and controlled in Matlab 2020a (MathWorks, Natick, MA, USA) and using Psychtoolbox (Brainard, 1997; Kleiner et al., 2007; Pelli, 1997). Gaze was recorded from one eye with a desk-mounted eye tracker (EyeLink 1000 Plus; SR Research, Kanata, ON, Canada) at a sampling frequency of 1000 Hz. To ensure accurate recordings before each block, a nine-point calibration was performed, and additional drift corrections were used at the start of each trial, where participants had to fixate on a fixation cross and start the trial by pressing the space bar. This allowed participants to take small breaks between the trials and start the trials in a self-paced manner.

### Stimuli

We used Blender 4.32 (Stichting Blender Foundation, Amsterdam, the Netherlands), an open-source 3D computer graphics application, to create 2 versions of a scene in which a spherical object rolls down a plank and falls into a container filled with liquid. In the first version, the object floats on the water surface as a plastic object with lower density would do; in the second version, the object drops to the bottom of the container as a metal object with higher density would do. Then, we created Consistent and Inconsistent stimuli by rendering the spherical objects in both scenes with a range of plastic and metal materials, approximation the appearances of (1) a beach ball, (2) bubble wrap, (3) yellow plastic, (4) rainbow plastic, (5) green plastic (plastic materials) and (1) iron, (2) rough copper, (3) hammered metal, (4) anti-slip metal, and (5) rusty iron (metal materials) (**Fig. 1b**). Materials for all scene objects, including container and liquid, were either available via the open BlenderKit add-on (royalty-free) or created in Blender. Environmental illumination was based on an HDRI lightmap (*Spiaggia di Mondello* by Poly Haven, from BlenderKit). All scenes were rendered using Blender’s Eevee engine at a resolution of 1920 × 1080 pixels with a total of 100 image frames. The animations were created at 30 Hz, resulting in 3.33 s per video. Finally, note that we did not create simulations or pick materials to be accurate in terms of real physics, but for generating plausible visual animations (similar to a computer game). All animations are available at: https://osf.io/e39hp/.

As all used materials were royalty-free, we don’t really need to include this table, whatever we prefer:

**Table 1.**
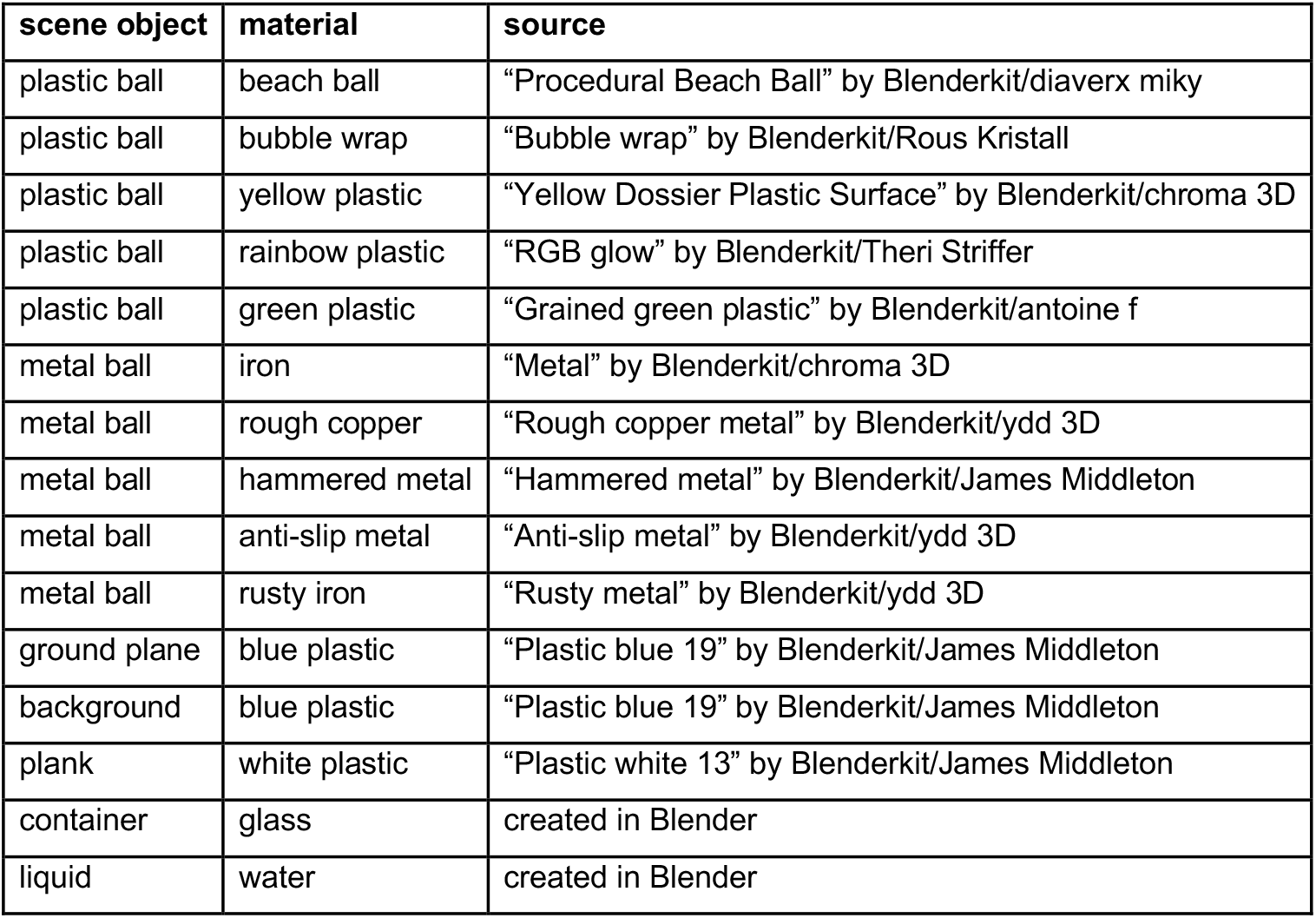
Blender materials and sources.

### Experimental conditions & paradigm

Participants completed two different tasks: a perceptual task and an interception task. The behavioral judgements and the eye tracking data is available at: https://osf.io/e39hp. All participants started with the Perceptual Task.

### Perceptual Task

The perceptual task was used to get an initial estimate of how participants judged the different materials. Each trial started by a button press, after which the first frame of the animation was shown for a random time between 1000 and 1500 ms. Then the animation started playing and stopped at the frame right before the object touched the water. Afterwards, participants used the keyboard to make a binary judgement about whether the object would sink or float. We showed each of the ten different animations (five plastic material variations and five metallic material variations) five times in randomized order (2 materials * 5 variations * 5 repetitions), so that each participant completed 50 trials in total.

### Interception Task

In our main experiment, participants saw the full animations and performed a dummy interception task. For that, we defined an area by two curved lines (see **Fig. 1a**), and asked participants to press a button when the object was in that area. The area was chosen so that the correct time window was similar for floating (Frames 69-96) and sinking (Frames 63-81) objects. This task was introduced to emphasize predictive eye movement behavior — note that our main analysis window was before or at water contact and thus well before the object entered the defined area. Critically, in the interception task we had three main conditions in separate blocks of trials: *Consistent, Inconsistent*, and *Mask*. In the *Consistent* condition the original animations were shown, where in all trials objects behaved in accordance with their material properties: plastic objects floated and metallic objects sank. In the *Inconsistent* condition, the behavior of the objects was also completely predictable (the same in all trials) but inverted: plastic materials sank and metallic materials floated. Finally, in the *Mask* condition, objects were masked by a black disc to remove all visual material cues, with half of the trials showing floating and half showing sinking objects.

Each trial was started by looking at a central fixation cross (also used for drift correction) and pressing the space bar, after which the static first frame of the upcoming animation was shown for a random time between 1000 and 1500 ms. Then the animation played until the end. The target area was visible at all times and participants were asked to press a button when the object was in the target area. The participants received feedback for correct responses by a thin green frame flashing around the animation. After the end of the animation, the next trial started by presenting the central fixation cross.

Each participant completed all three conditions in separate blocks of trials. To study intuitive physics (anticipation of realistic behavior without previous training), half of the participants completed the order *Consistent* – *Inconsistent* – *Mask*, and the other half *Inconsistent* – *Consistent* – *Mask*. The *Consistent* and *Inconsistent* conditions had 70 trials (2 materials * 5 variations * 7 repetitions) and the *Mask* condition 40 trials (2 materials * 20 repetitions), so that each participant completed 180 trials in total. Within each condition, trials were presented in randomized order.

### Data Analysis

Eye movement data were digitized online and analyzed using custom Matlab code. The code is available at: https://github.com/AlexanderGoettker/MaterialAnticipation.

First, blinks were linearly interpolated, and the eye position was filtered with a second-order Butterworth filter, with a cutoff frequency of 30 Hz. Then for each trial we looked at the vertical position aligned to the moment the object touched the water. To visualize the data we saved the vector from 800 ms before to 800 ms after water contact. To get a single metric, we also estimated the vertical eye position at water contact as the median of the vertical position between 50 ms before and 50 ms after water contact.

Our main analysis is based on a decoding approach, where we trained a classifier to distinguish between floating and sinking objects based on the vertical eye position. For that we took each time point in the aligned vertical eye position vectors and got these data across for trials per condition. We then randomly selected 25% of trials as test and the other 75% as training data for a linear classifier (‘classify’ in MATLAB). We repeated this procedure 50 times, and computed for each timepoint the average proportion correct across these iterations. Since we were running the classifier in 1-ms steps, we used a moving average with a window of 10 ms to smooth the classification performance. This provides an estimate of the time course of predictive eye movement behavior. To express this in a single metric, we also computed the median of the proportion correct estimates between 50 ms before and 50 ms after water contact.

To analyze the effect of uncertainty about the material properties, we looked at all possible pairs of plastic and metallic stimuli. We then computed the average proportion correct for each pair in the perceptual task and compared it to the average vertical distance in the eye position for both stimuli at water contact. If the predictive effect scales with uncertainty, the difference in eye position should be larger for pairs that were more accurately identified in the perceptual task.

To estimate the changes in predictive eye movement behavior within a block of trials, we used a similar approach. Instead of running the classifier across time relative to the moment of water contact, we only used the trial estimates of the vertical eye position at water contact. Half of the trials were used as training data, again across 50 iterations, to classify each individual valid trial of a participant to estimate proportion correct over time in the block. To estimate performance, we again averaged across all 50 iterations and smoothed the estimate with a moving average of seven trials. Importantly, to not bias performance early in the block, we used an asymmetric moving window of always the last seven trials that were available. For the early trials, the average was therefore based on fewer trials (e.g., on 2 trials for the second trial). To make the time course comparable between conditions and participants with different amounts of valid trials, all vectors were temporally normalized to range from 0 to 100% of the valid trials in a block. To estimate the learning over time, we averaged these time-course data across subjects. To quantify the shape of the learning, we fitted a simple exponential learning function with three free parameters to the data

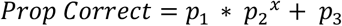

with *x* as the time in the block, *p*_*1*_ and *p*_*2*_ describing the learning rate and *p*_*3*_ being the asymptote of the curve. We estimated the stability of the parameters by bootstrapping. We used 100 iterations where we randomly only selected 25 participants and then fitted the curve. This allowed us to estimate the CI of the parameter estimates. To study intuitive performance (without training), we also extracted the classification performance for the very first trial.

### Exclusion Criteria

In the perceptual task we used all trials for the analysis. In the interception task, we removed individual trials from the analysis when there was either (i) missing data for more than 500 ms, or (ii) unclear missing data that were not labelled as blinks by the Eyelink eye tracker. Based on these criteria, we included 5129 out of 5520 (95%) for the analysis.

